# Mechanical Force Works as a Biological Cue in Postnatal Murine Tendon Development

**DOI:** 10.1101/2023.06.02.543346

**Authors:** Yuna Usami, Xi Jiang, Nathaniel A Dyment, Takanori Kokubun

## Abstract

The musculoskeletal system provides structural stability and coordination to enable movement. Tendons have the essential role of efficiently transmitting force generated from muscle contraction to bone to enable ambulation. In doing so, they resist high external forces. In fact, muscle contraction during embryonic development is required to maintain tendon growth and differentiation. Nonetheless, defining the types and magnitudes of loads that act on tendons during embryonic and early postnatal periods is quite difficult. In this study, we aimed to define the physiologic limb movement and forces experienced during these stages in the murine model. We found that late-stage embryos had limited amniotic space, which attenuated limb movement. In the neonatal phase, physical ability, as measured by rollover function and locomotion, increased. These changes, which likely corresponded to increased forces applied to the tendons, corresponded with the expression of tenogenic markers during the embryo to postnatal phase. In particular, we found that the upregulation of *Scx* and *Tnmd* correlated with increased movement during the two weeks after birth. Our results help define the spatiotemporal role of mechanical force, including internal and external factors, in tendon growth and development.

**Highlights:**

1. Assessed limb movement in amnion.
2. Space limitation attenuated limb movement in the late-stage embryos.
3. Defined the mechanical force from the limb’s physiological environment.
4. Scx and Tnmd were upregulated synchronically with rollover function and locomotion.
5. Mechanical forces may work as the cue of tendon development

## 1. Introduction

The musculoskeletal system provides structural stability and the ability for physical movement. Not only do these tissues transmit mechanical forces to enable movement, but they require loading to properly develop and grow. In fact, neural fibers connect to the muscle around E12.5 (Martin, 1990; Suzue, 1996), and skeletal muscle can begin to generate contractile force around Embryonic day (E) 13.5 of the mouse(Michael R. Carry et al., 1983). Then, tendons transmit the force that the muscle generates to the skeleton to facilitate limb movement.

The tendon is a dense fibrous connective tissue that can resist repeated mechanical loading from muscle contraction and other external forces. There are key molecular signaling processes that drive tendon development (Asahara et al., 2017; Subramanian and Schilling, 2015). For instance, critical signaling pathways such as transforming growth factor-beta (TGF-b) (Pryce et al., 2009), Mitogen-Activated Protein Kinase (MAPK) (Schwartz et al., 2015), and hedgehog signaling regulate the formation of the tendon mid-substance and/or enthesis (Fang et al., 2022). In addition, key transcription factors, such as Scleraxis (Scx) (R Schweitzer et al., 2001), mohawk homeobox (Mkx) (Ito et al., 2010), and early growth response Egr1/2 (Lejard et al., 2011), and SRY-box transcription factor 9 (Sox9) (Blitz et al., 2013; Sugimoto et al., 2013) drive differentiation and maturation of tendon cells during embryonic and early postnatal development. While these biological cues are well recognized in promoting tenogenesis, there is a limited understanding of how mechanical cues contribute to tenogenesis during this critical period.

During embryonic development, musculoskeletal tissues are exposed to several types of mechanical forces. For instance, muscle contraction is important for the continued growth of tendons (Huang et al., 2015) but is not critical to the specification of tendon cells. Elongation of the skeleton also imparts growth-mediated stresses and strains on the tendon to drive continued differentiation and maturation (Schiele et al., 2013). These forces imparted on the limb are essential for the proper development of the limbs, with limited limb movement from various factors resulting in fetal akinesia. Prenatal limb movement is affected by the mam’s physical movement and the resistance force with amniotic fluid in the amnion, like a subaqueous condition. In contrast, in the neonatal phase, newborn pups are exposed to a completely different environment from in the amnion. They have to move their limbs against gravity. Then, they acquire enough muscle contractile function to move their limbs freely. The central nervous system gradually matures at that time with postural reflexes starting to appear (Feather-Schussler and Ferguson, 2016; Semple et al., 2013). These physical movements, in addition to bearing weight during ambulation, increase the loads being applied to the limbs. However, the direct relationship between physical movement and tendon development during this period is not fully understood.

In this context, we hypothesized that the mechanical force generated by muscle contraction and external force depending on the physical behavior on the limb have independent effects as a mechanical cue on tendon development, leading to mechanobiological phenomena. To address this hypothesis, we investigated the changes in limb movement during the embryonic to the early postnatal period that may impact the mechanical loads applied to the tendon. Additionally, we also explored the relationship between the changing mechanical force and the expression of tenogenic factors and changing tendon morphology.

## 2. Materials and methods

### 2.1. Mice

All animal procedures were approved by the Animal Research Committee and were consistent with animal care guidelines (Approval Number: 2019-10). The mouse lines were used wild-type C57BL/6J (Tokyo Laboratory Animals Science Co., Ltd) backgrounds were bred in-house. For embryo harvest, timed mating was set up in the afternoon, and identification of a mucosal plug the next morning was considered 0.5 days of gestation (E0.5). We used in the study were from littermate mice. Animal subjects were housed in a controlled environment with a 12:12-h light-dark cycle with ad libitum access to water and food. The experimental design is illustrated in Fig. 1.

**Fig. 1.**
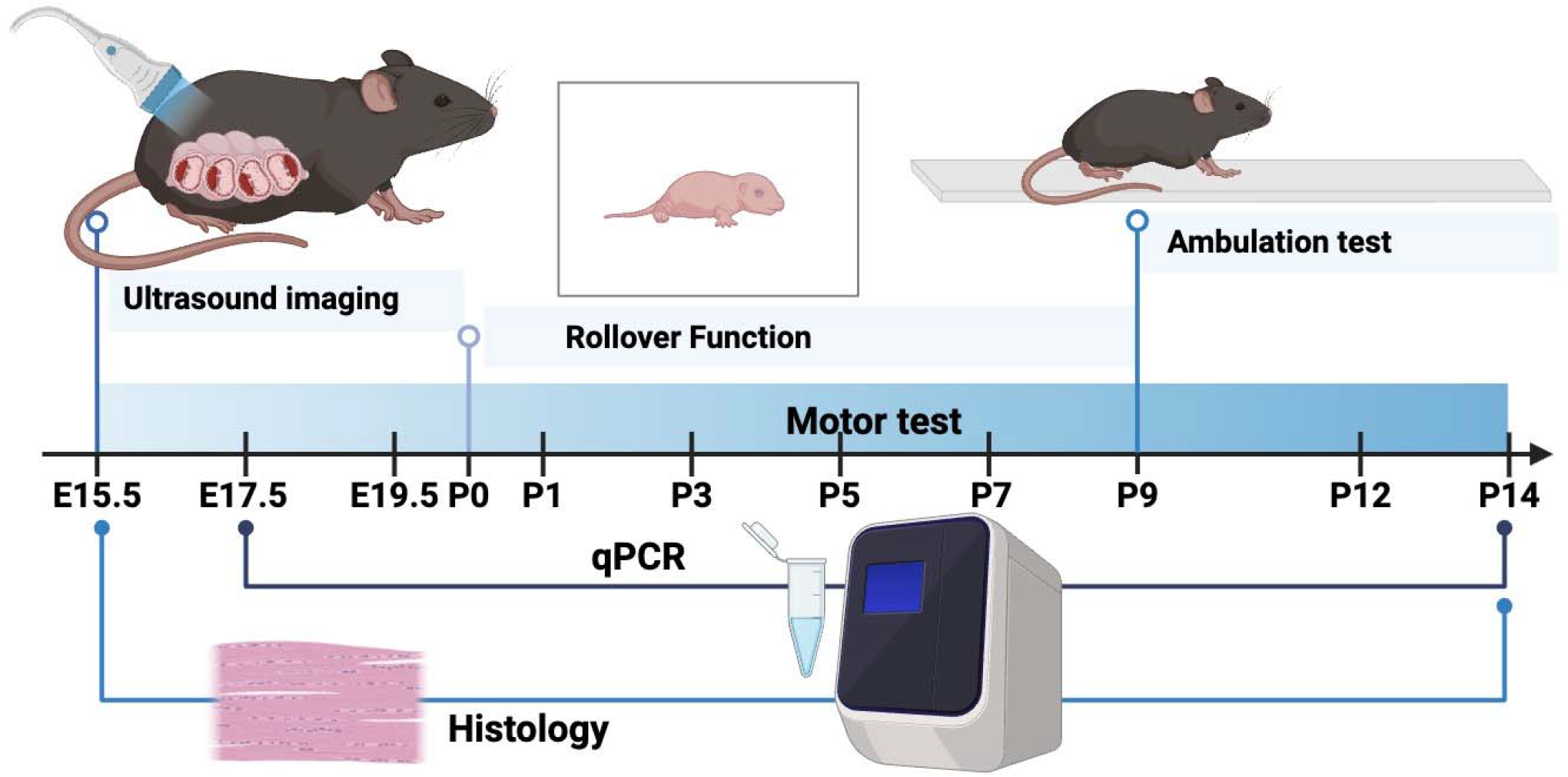
Experimental scheme of this study. Research Design Overview

### 2.2. Ultrasound imaging

The embryonic movement was observed using Real-time Ultrasound Imaging System (Vevo3100, FUJIFILM VisualSonics, Inc., Toronto, Canada). This investigation focused on to reveal the actual embryonic limb movement in the amnion at the E15.5, E17.5, and E19.5 in mice. Pregnant mice were transferred to the imaging center. Ultrasound imaging was performed under general anesthesia, induced with 5% isoflurane and maintained with isoflurane 2%. During the scan, physiological changes were monitored with in-built ECG electrodes, and body temperature was maintained at 37 °C. Their abdomen’s fur was removed using a chemical hair remover, and the pre-warmed gel was applied to their abdominal region. To test the effect of anesthesia, we performed the same protocol under the non-anesthesia condition as well. Two days before imaging, we removed the hair in the abdominal region of pregnant mice under the isoflurane. At the observation, we carefully dislocated the cervical of the mom mice and quickly observed the limb movement of the embryo using ultrasound imaging at the E17.5. We tracked the embryo’s limb movement, heartbeat, and blood flow in the embryo’s limb to confirm its condition. The mom’s body temperature was maintained at 37 °C during the ultrasound scan.

### 2.3. Neonatal Motor Tests

Mouse pups were tested for motor behavioral development according to the previous study (Feather-Schussler and Ferguson, 2016) to evaluate the performance of physical movement. Pups were tested 6hr (±1.5hr) after the light cycle started to eliminate the time-of-day difference in behavior. Pups were moved from the dam for no more than 15 minutes to avoid losing their body heat, hunger, and separation issues. The rollover function is commonly used as the motor ability test for mice pups to be able to flip onto their feet from a supine position. In accordance with a previous study (Feather-Schussler and Ferguson, 2016), mice aged P0 to P7 tested the rollover function. Ambulation was assessed using the following scale: 0 = no movement, 1= Crawling with asymmetric limb movement, 2 = slow crawling but symmetric limb movement, and 3 = fast crawling/walking. In detail, the head and tail of a crawling mouse were low to the ground, and the head began to rise during the transition from crawling to walking. The transition was complete when the head and tail were elevated, and only the front of the hind paw touched the ground. The original test arrowed mice to score ambulation arrowed be repeated many times. Still, this research arrowed them for 1 minute considering the effect of fatigue. The score of each mouse was adopted at the highest score three times based on the mentioned ambulation, was no potential for learning during the test.

### 2.4. Histology

After sacrifice, forelimbs were skinned (E17.5∼) and fixed in 4% paraformaldehyde for 24 hrs. at 4 □. Postnatal tissues were decalcified in 50 mM EDTA for one day to 2 weeks at 4 □, then incubated in 5% sucrose (4hr) and 30% sucrose (overnight) at 4 □. Limbs were then embedded in an OCT compound, and 12 μm longitudinal cryosections were collected using Cryostar (Thermo Fisher). Sections were stained with Alcian blue and Hematoxylin/Eosin. For immunostaining, sectioned slides were rinsed in PBS-T and blocked in Dako Antibody Diluent with Background Reducing Components (Agilent Technologies) for 1h at room temperature using M.O.M. kit (Vector Laboratories). Then sections were incubated in primary antibodies Collagen type□ (Col1) (Abcam) overnight at 4□, and secondary antibodies for 4h each at room temperature, and My32 (Sigma) was used to detect muscle-specific type II myosin heavy chain (MHC) for 30min at room temperature with secondary antibody Dylight588 (Abcam) and Cy3 secondary detection (Jackson ImmunoResearch) and remove unwanted fluorescence (Vector TrueVIEW Autofluorescence Quenching Kit with DAPI, Vector). DAPI was counterstaining to visualize cell nuclei. Sections were observed under a BZ-X710 microscope (Keyence).

### 2.5. In situ hybridization (RNA scope)

Lower limbs were fixed in 10% neutral buffered formalin for 2 days, transferred to 30 % sucrose in PBS overnight, and embedded in OCT compound (Thermo Fisher). The ankle was cut in the sagittal plane to investigate the Achilles tendon. All sections were made from undecalcified joints using cryofilm (Dyment et al., 2016). The filmed sections were glued to microscope slides using a chitosan adhesive and rehydrated prior to imaging. RNAscope in situ hybridization (ISH) multiplex version was performed using a probe targeting Scleraxis (439981-C3) according to the manufacturer’s protocol (Advanced Cell Diagnostics).

### 2.6. RNA isolation, reverse transcription, and qRT-PCR

Total RNAs were extracted from the E17 to P14 Achilles tendon using ISOGEN (Nippon gene). The reverse transcription was TaqMan™ Gene Expression Master Mix. Primer sequences used are *Scx, Transcription growth factor-*β*2*(*Tgfb2*), and *Tnmd*. For mRNA level analyses, gene expression changes were quantified using the -2ΔΔCt method. Housekeeping was *glyceraldehyde-3-phosphate dehydrogenase* (*Gapdh*).

### 2.7. Statistical analysis

GraphPad Prism (version 9.4.1) was used for statistical analysis and to generate all graphs. Non-statistically significant comparisons were not shown. A one-way ANOVA with Tukey-Kramer posthoc test was performed across ages for each measured parameter.

## 3. Results

### 3.1. Spatiotemporal changes in limb movement during late embryonic and early postnatal growth

We observed embryonic limb movement by using ultrasound imaging. In embryos from E15.5 to E19.5, we did not detect spontaneous limb movement of fore/hind limbs except independent of their heartbeat (Fig. 2, Supplementary Video 1, 2). Depending on the pushing force from the ultrasound probe, they showed whole-body rotation slowly in the amnion. However, The space in each amnion decreased with age the size of the embryo increased, which limited limb movement in the uterus. As a result, we did not detect voluntary limb movement in the late embryonic phase (Fig. 2D). Additionally, we confirmed the effect of anesthesia for the embryo’s limb movement (Supplementary Video 3), but there were no differences on anesthesia condition.

**Fig. 2.**
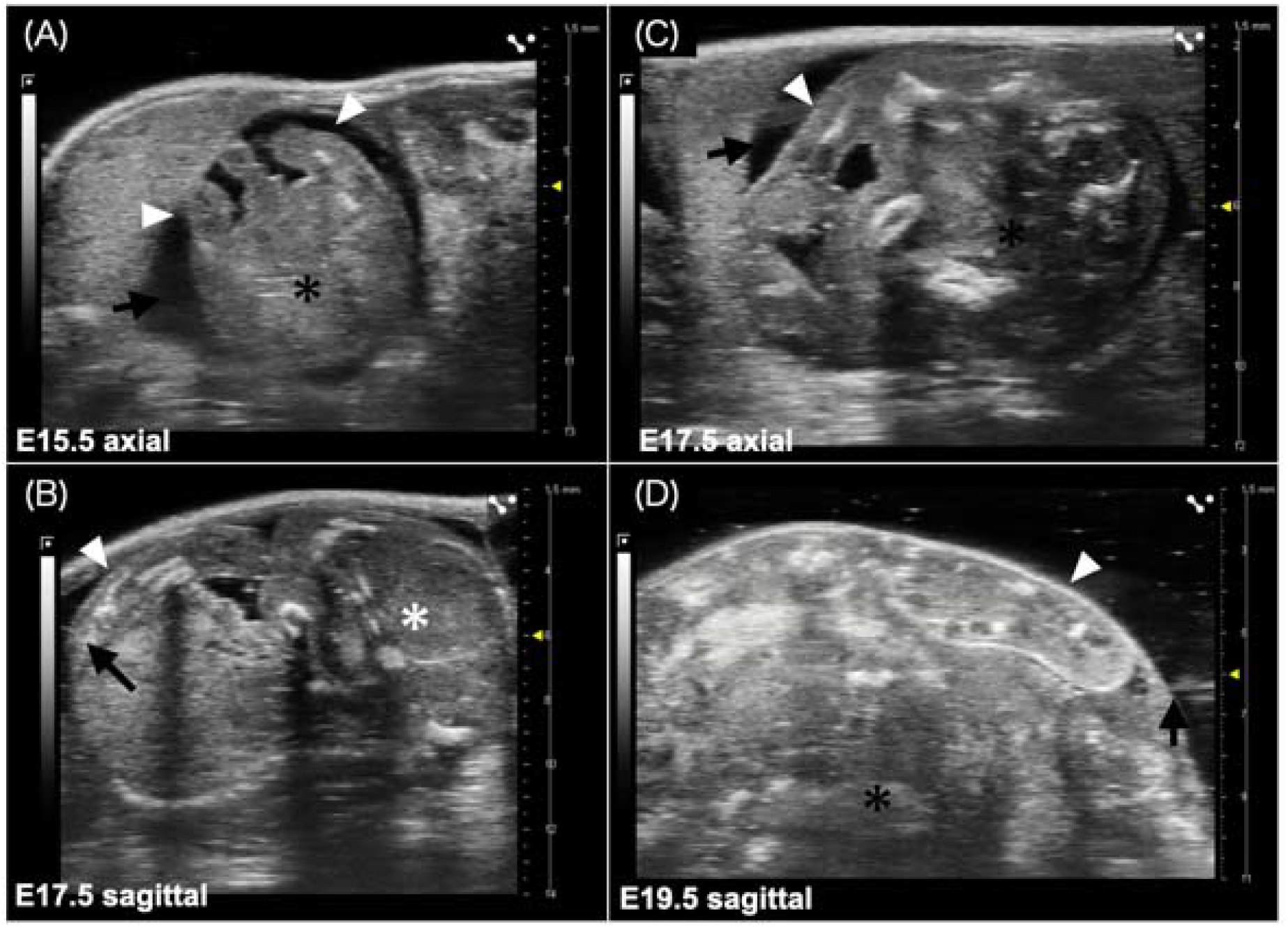
Ultrasound imaging showed the embryo limbs had no detectable limb movements. The axial shows limbs of (A) E15.5 and (C) E17.5 and sagittal imaging of (B) E17.5 and (D) E19.5 embryos. E19.5 had markedly reduced space compared to E15.5 and E17.5. White arrowheads denote hindlimb and black arrows mark the amniotic space in each embryo. The trunk and the head were marked with asterisks.

### 3.2. Changes in limb movement and gait pattern with postnatal age

The body weight increased gradually with age in the postnatal phase (Fig. 3A). P0 mice were lying only in a supine position (i.e., non weight-bearing), and their limb movement was minimal. At P1, they irregularly moved their fore and hind limbs in a supine position. From P1 to P5, they gradually were able to flip onto their limbs from a supine position. All mice were able to turn over by P7 (Fig. 3B), indicating that weight-bearing likely increased substantially after this age. This age also corresponded to when the pups started to ambulate and ambulation improved from P9 onwards (Fig. 3C). The gait of P9 showed crawling with asymmetric limb movement. With increasing gait scores, they obtained a symmetric gait pattern with age from P9 to P14. In fact, their tail and hock became elevated during ambulation at P14 (Fig. 3D).

**Fig. 3.**
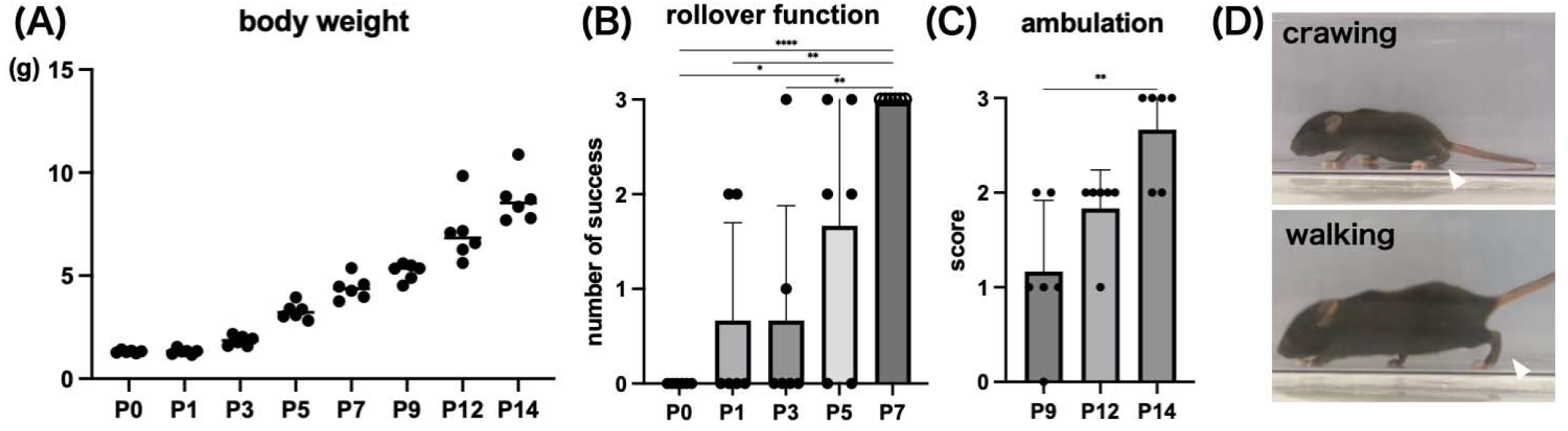
Neonatal motor tests demonstrated increased ambulation and limb movement during early postnatal growth. (A) Bodyweight increased with age. (B) The number of successes in rollover function test increased with age (3 tests per mouse) [P0 vs. P5 P=0.0367, P0 vs. P7 P<0.001, P1 vs. P7 P=0.0019, P3 vs. P7 P=0.0019]. (C) The ambulation score (0 = no movement, 1= Crawling with asymmetric limb movement, 2 = slow crawling but symmetric limb movement, and 3 = fast crawling/walking) was adopted at the highest score of three repeats [P9 vs. P14 P=0.0012]. Error bars represent SDs. (D) The hock (white arrowhead) began to rise from crawling to walking, as agreed with the previous study (Feather-Schussler and Ferguson, 2016). (n = 6/time point)

### 3.3. Gene expression changes with late embryonic to early postnatal growth

We performed qPCR to understand the relationship of tenogenic gene expression with changing mechanical force associated with the physical environment or locomotion. However, we could not analyze the tissue of E15 because it was too immature to collect muscle and tendon tissue under the stereoscopic microscope. We found the expression of each tenogenic gene but no significant differences in expression levels in each time point. However, the expression levels of Scx gradually increased for P12, and the expression level of Scx displayed a higher level before P5 compared to E17.5 to P3, Tnmd as well. The *TGFb2* showed the highest expression of P5. The values of all time points were stable at the standard value of E17.5 (Fig. 4).

**Fig. 4.**
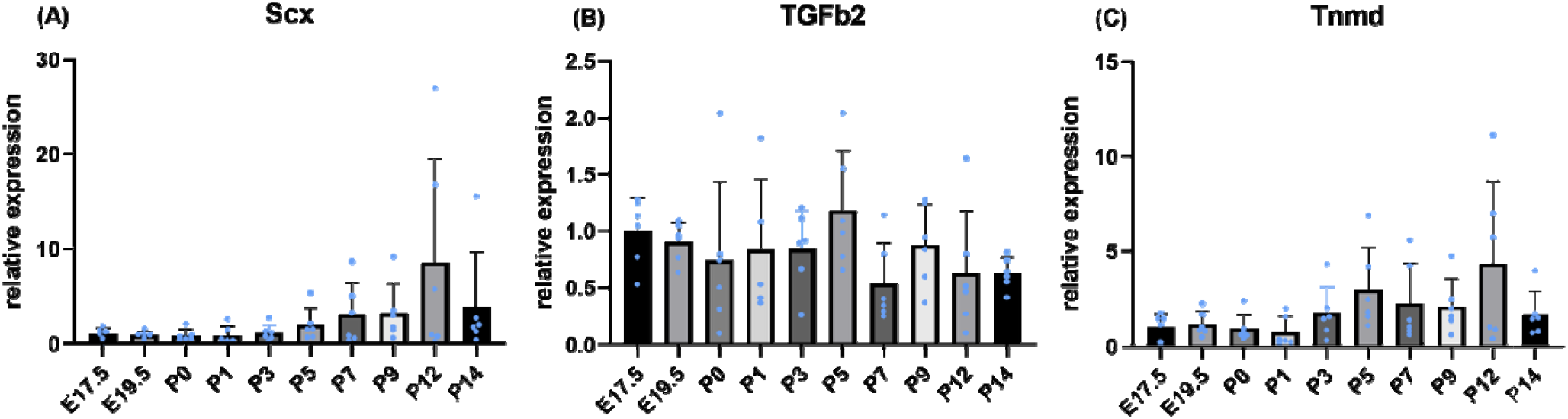
Gene expression changes during late embryonic and early postnatal growth. Results of mRNA expression. Gene expression analysis in Achilles tendons of mice. (A–C) Quantitative analysis of qPCR for *Scx, TGFb2*, and *Tnmd* expression. Target gene expression values were normalized to the stable housekeeping gene Gapdh, and then relative to E17.5 expression levels. Values are mean ± SD, n = 5-6 for each group. Difference between groups was tested using ANOVA and the Tukey-Kramer host-hoc test. There were no significant differences.

We also conducted RNAscope RNA FISH for Scx to visualize the spatiotemporal changes of these key tenogenic transcription factor with age. We found clear expression in the midsubstance of developing tendon throughout the time period. The size of tendon and bone was increasing over this time period as well. The shape of the cells nuclei gradually elongated in the early postnatal phase (P0 to P7, Fig.5), and the number of tendon cells was reduced in the mid to late postnatal phase (P9 to P14, Fig.5).

**Fig. 5.**
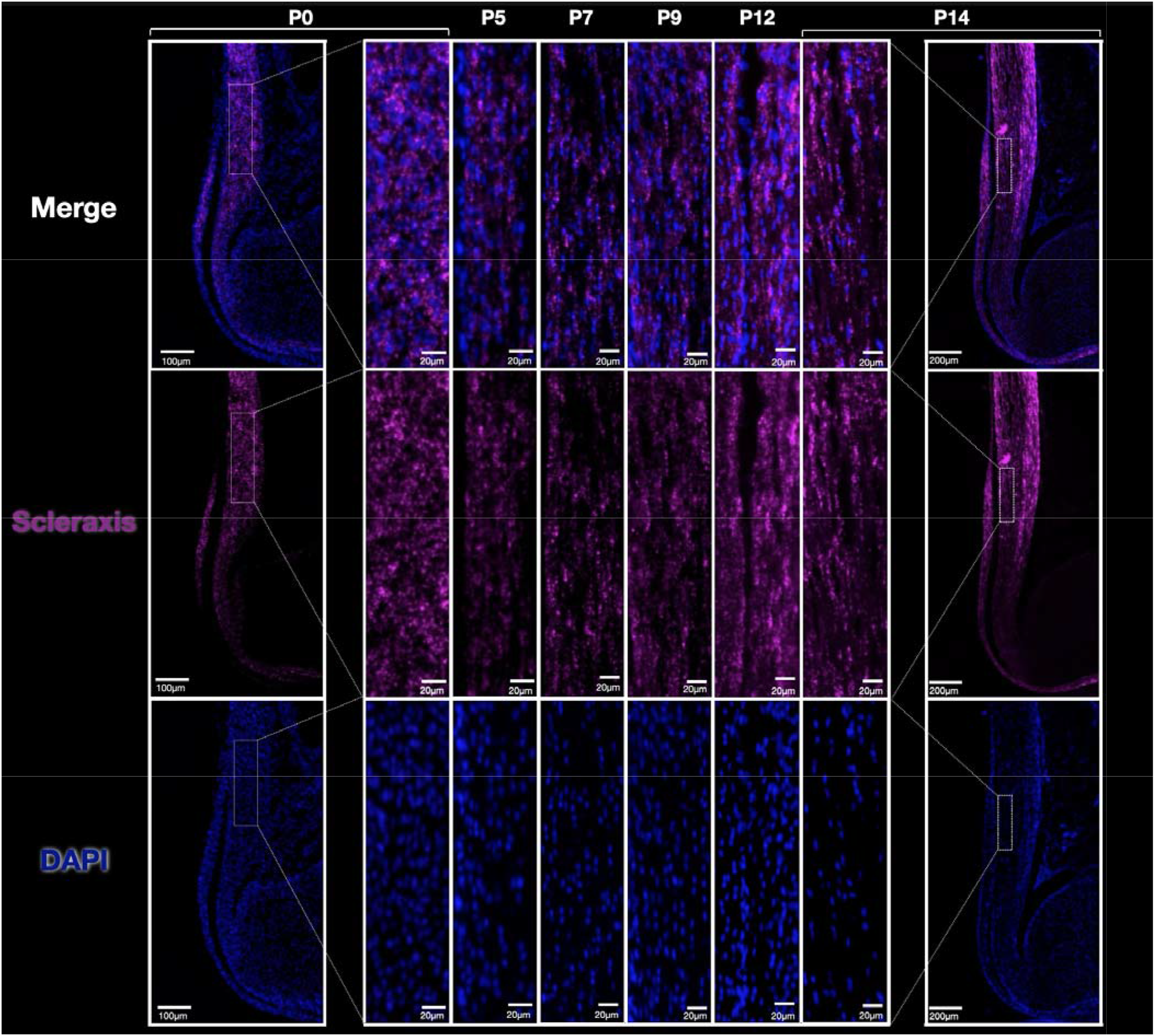
Scx expression via *in situ hybridization* was consistent during early postnatal growth. RNAscope showed a clear expression in the midportion of the developmental tendon through the investigated time period. The size of the tendon and bone increased over time, as expected. The nuclear shape of tendon cells was gradually elongated in the early postnatal phase (P0 to P7), and the number of tendon cells was reduced in the mid to late postnatal phase (P9 to P14). These changes occurred concurrently with increasing the mechanical force corresponding with physical movement.

### 3.4. ECM production and organization increases with age

In Alcian Blue and HE stains, the length of the tendon became longer with time course, and the line of the muscle-tendon junction became remarkable. At E15.5, the cohesion of cells was observed at the site where tendon tissue will be formed, but no extracellular matrix (ECM) stained with eosin was observed at this time. At E17.5, the cell density increased, and the border between the tendon tissue and the surrounding tissue became clear. It was a simple cell aggregation image, but in the tissue, after P0, the tendon cells became elliptic and aligned in the long-axis direction (Fig. 6A). We detected localization of collagen type 1 (col1), the most abundant form of collagen in tendons, in immunofluorescence staining. We confirmed collagen type 1 after E17.5 and increased for P14 (Fig. 6B). At P14, fibril formation progressed with the increased ECM in the long-axis direction, and the decrease of nuclei was confirmed. Like tendons, the muscle tissue had cell aggregation during the embryonic period, but at P3, the muscle-tendon junction became apparent, and the muscle tissue fiber arrangement was formed (Fig. 6A).

**Fig. 6.**
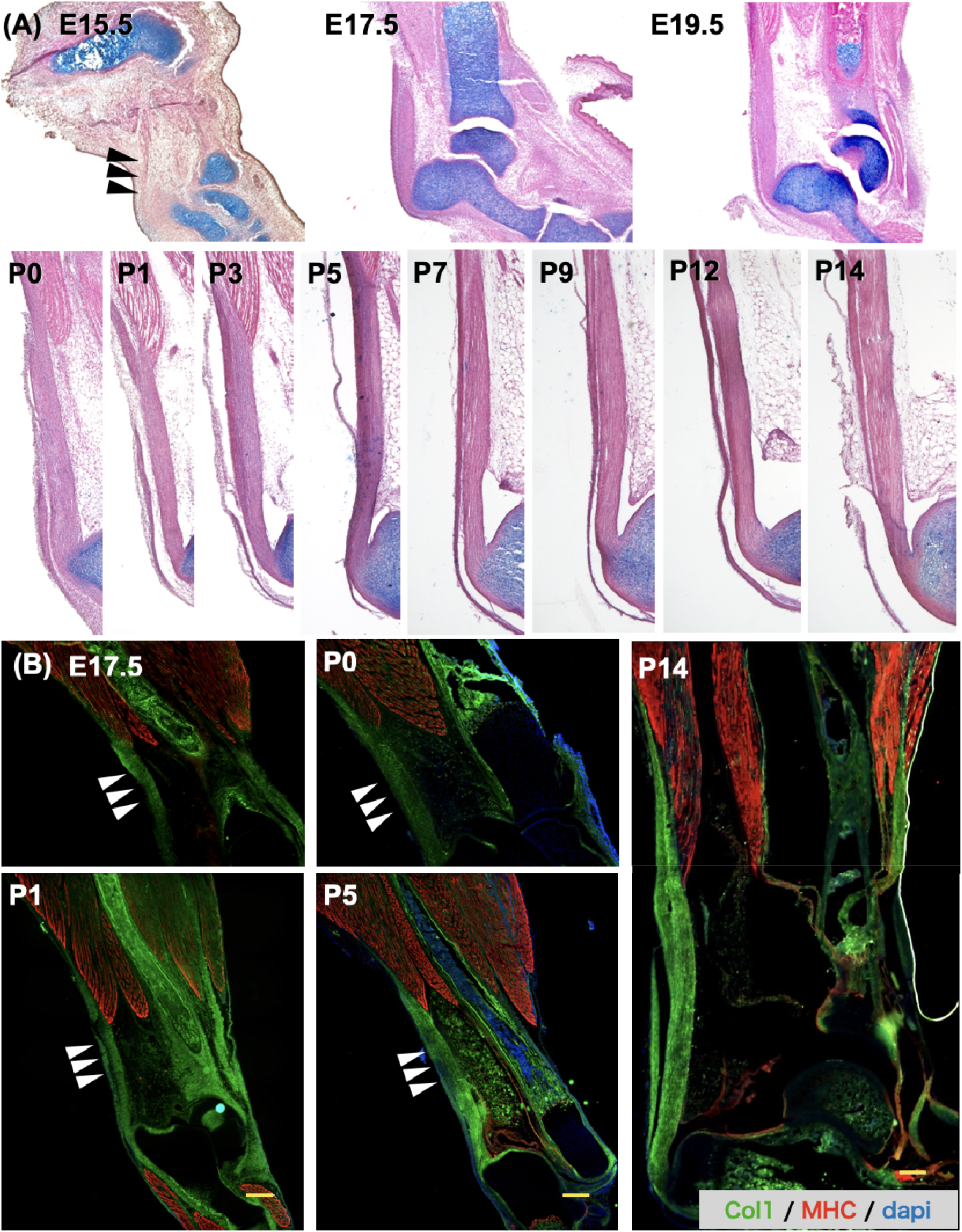
Alcian blue and HE stains, and Immunofluorescence of hindlimbs showed increased Achilles tendon ECM. (A) Sagittal sections of the ankle were shown. The tendon matured gradually with the time course. The arrowhead shows the Achilles tendon (E15.5). (B) Double immunostaining of Col1 (green) and muscle (MHC) was performed on frozen sections. Scale bar = 200µm.

## 4. Discussion

Tendons are strong and durable tissues that are resistant to high tensile forces. During growth and development, they experience mechanical force from muscle contraction, growth mediated stresses and strains, and weight bearing. The magnitude of these forces are dynamically increasing through the embryo to the early postnatal phase. The relationship between the tenogenic differentiation process and increasing mechanical force within this tissue is poorly understood. Therefore, in this study, we tracked limb movement in utero using an ultrasound imaging technique, passive joint motion with physical activity and weight bearing by P7, and muscle and passive forces during ambulation. We also analyzed the growth process of the murine Achilles tendon from its late embryonic stage through its first two weeks of postnatal growth. During this phase, mice gain neuro-muscular function, resulting in limb movement, ambulation, and weight-bearing. Surprisingly, we could not detect limb movement during the late embryo phase using our high-resolution ultrasound imaging. In contrast, after birth, pups moved their limbs in a supine position at first. Then, their limbs were exposed to higher mechanical force, as they began to bear weight. Through this phase, Scx gene expression noticeably increased concurrently with changing and increasing the physiological movement. The mechanical force with physiological movement is likely critical in the tendon maturation process during the early postnatal phase in the murine.

### 4.1 The Importance of Limb Movement in the Late Embryo Phase to Tendon Maturation in Murine

We observed embryonic movement in the uterus using high-resolution ultrasound imaging. Previous works from neurological research showed that the first innervation of the mouse limb appeared after E12.5 (Martin, 1990; Michael R. Carry et al., 1983). Mouse embryos at the early developmental stage of E12.5 can generate organized limb movements in the ex vivo transplacental perfusion method. Suzue et al. detected mouse hindlimb movement in an area of 300μm square, which is insensible movement (Suzue, 1996). Lipp et al. exposed freshly explanted limbs to acetylcholine to determine when muscle contraction begins. They reported normal mice limb muscles at E13.5 and E14.5 had confirmed contractile properties but not at E12.5 (Lipp et al., 2023). These previous studies revealed that murine limb muscles gain neurological innervation at E12.5 and contractile properties at approximately E13.5. Referencing these papers and using transgenic mice with reduced muscle contraction, previous papers mentioned the muscle contraction is needed for tendon elongation during the embryo phase in mice (Huang et al., 2015; Kahn et al., 2009; Tsinman et al., 2023). These data suggest the importance of muscle contraction in tendon development after the E15.5 late embryo phase, but there are no studies that tracked actual limb movement in the uterus and amnion. Here, we tracked limb movement at E15.5, E17.5, and E19.5, but surprisingly, we did not detect spontaneous limb movement in the uterus and amnion during the late embryo phase by high-resolution ultrasound imaging (Supplementary Video 1-3).

The embryo lives in the small space of an amniotic membrane, so the body gradually arches, and limbs are folded in front of the trunk depending on the skeletal maturation. Physiological development led to muscle maturation as well, but dramatically reduced space in the amniotic membrane and disturbed embryonic limb movement. This environment at the later embryonic stage is similar to a chick embryo. Bradley et al. reported the chick embryo limbs within the egg maintained a flexed position, and movement was potentially constrained by the shell wall (Bradley et al., 2014). Additionally, the number of offspring may further limit space for limb movement in the uterus, especially in mice with larger litter sizes. The same effect was reported, which is that the number of offspring affects limb movement in human fetuses. Comparisons of fetal movements between singleton and twin fetuses using a 4-D ultrasound image revealed that the frequencies of any kind of movements in twin fetuses were significantly lower than those in singleton fetuses (AboEllail et al., 2018). There could be multiple reasons that explain the discrepancy between the lack of limb motion detected in our study and the tendon phenotype found in mice that lack skeletal muscle contraction (Huang et al., 2015; Tsinman et al., 2023). For instance, our ultrasound technique may not be sensitive enough to detect the minor limb movement that may be required for proper growth. However, limited limb movement in utero does not negate the ability of the muscles to undergo isometric contraction, which may be enough for continued tendon growth and maturation. Concentric and eccentric contractile forces are likely to increase during the postnatal period as the animal begins to ambulate, which may further contribute to the substantial growth via matrix accrual during this period. In bone development, Nowlan et al. reported movements of the mother or littermates to compensate for passive stimuli during embryonic skeletogenesis, using a murine model lacking skeletal muscle (Nowlan et al., 2012). Previous studies in neurological research reported that late pregnant mothers are active throughout late gestation, similar to nonpregnant females. Locomotor behaviors in pregnant mothers are rearing, head and body grooming, and stirring. That provides a sensory experience for rodent pups in utero (Ronca et al., 1993). In mammals, these mother’s movements have the potential to provide mechanical force for the embryo to make joint movements by touching limbs to the amniotic membrane. Although clear limb movement in mice in utero has not been identified, we cannot deny that muscle contraction contributes to tendon development. Our study offers additional evidence on the types and magnitudes of forces that may be required.

### 4.2. Mechanical Force in Late Embryo to Postnatal Phase Related to the Tendon Development

After birth, the limbs are released from space limitation and begin dynamic movement in the supine position. These joint movements are generated by the force of muscle contraction against gravity. We expect a change in the mechanical environment for limbs to allow free movement; this leads to increasing muscle activity. However, P0 mice, immediately after birth, did not have enough matured muscle function to move their limbs. As a result, the limbs had only limited movement because gravity was applied to their limbs even if they didn’t bear weight loading.

To confirm the biological tendon response to the gradually induced mechanical stress, we analyzed the expression of tendon development-related genes. Scx is known for specific tendon maturation markers from the early progenitor stage to the formation of mature tendons. Previous studies showed that Scx expressions reflect changing the mechanical signals in the mouse limbs (Eloy-Trinquet et al., 2009; Havis et al., 2016). This study focused on the different tendon developmental genes; however, finally, these genes activate the expression in developmental tendons. Notably, Maeda et al. showed mechanical forces maintain Scx expression through the TGF-β signaling pathway (Maeda et al., 2011). TGF-β signaling has a role in adapting to tension released in response to force. TGF-β acts on tenocytes to alter their morphology and ECM production, revealing a feedback mechanism (Maeda et al., 2011; Subramanian et al., 2018). Tnmd is a fine-tuner of tenocyte proliferation and collagen fibril maturation (Docheva et al., 2005) and requires tendon adaptation to mechanical load (Dex et al., 2017). Scx directly activates Tnmd to regulate tenocyte differentiation and maturation (Shukunami et al., 2018). In this context, we analyzed whether the expression of these genes is related to changing the mechanical force depending on motor behavior in tendon maturation.

During E17.5 to P1, tendon development-specific genes; *Scx, TGFb2*, and *Tnmd* have not shown dynamically changed. With the ultrasound imaging, the late embryo showed no specific limb movement in the amniotic membrane, and early postnatal could not move their limb immediately because of the gravity environment. The neonatal mice showed little movement of their limbs during P0 to P1. Concurrently, qPCR results showed the expression of *Scx* and *Tnmd* showed low-level expression during E17.5 to P1 (Fig.4). The environment of the fetus has changed dramatically in terms of circulatory, respiratory, nutrient, and mechanical conditions. Actually, P0 to P1 mice showed radically frequent limb movement. Nevertheless, the expression of tenogenic genes showed a slight change pre/post-delivery. Bone cells can receive passive stimuli, like compression, and share force with mom’s behavior directly. In contrast, tendon cells need joint movements to get the stretching force depending on the muscle contraction or passive stimuli. However, the space in the amniotic membrane is limited, so the amount of stretching force to the tendon cell may be quite small. Our results suggest that passive and external mechanical force may be a piece of essential factor in tendon development in the late embryo to early postnatal phase. Additionally, our results give another point of view to consider the results of transgenic mice studies because previous studies do not refer to this perspective (Usami et al., 2024). Further study is needed to reveal the role of these external mechanical stimuli on tendon development.

### 4.3. Tendon development in the Mid to Late Postnatal Phase

Next, we assessed the rollover function to define how much increase the passive mechanical force on the postnatal limb depending on the weight bearing with behavior patterns. Before getting the rollover function, muscles produce limb motion in the air against gravity. Then, when they gained the rollover function, their limbs touched the ground and were exposed to mechanical force by their own weight. Our results showed that P5 mice increased their time in a prone position, which led to increasing of mechanical force gradually on their limbs. After getting the rollover function completely at P7, limbs are exposed to high mechanical force depending on the weight-bearing load. In this phase, *Scx* and *Tnmd* tended to increase the expression, but there was no significant difference.

Next phase, they started to ambulate. At P7-9, mice showed crawling with asymmetric limb movement (Fig. 3, 7). This means these aged pups did not have enough muscle function to bear their body weight. At P10 to 12, their hock gradually changed to non-contact with the ground during the stance phase. After transitioning from crawling to walking at P14, they could raise the hock, which means maturing the muscle function (Fig. 7). In this phase, mechanical force on the tendon was maximumly increased because muscles were matured and generated enough force to bear their own weight. These forces clearly generated a high stretching force for the developmental tendon. According to the development of ambulation, *Scx* expression in qPCR tended to increase to P12. The results of ISH also showed the obvious *Scx* expression in the Achilles tendon through the postnatal phase. These results indicate that increasing mechanical force depending on the changing motor behavior probably has an important role in tendon development in the postnatal phase.

**Fig. 7.**
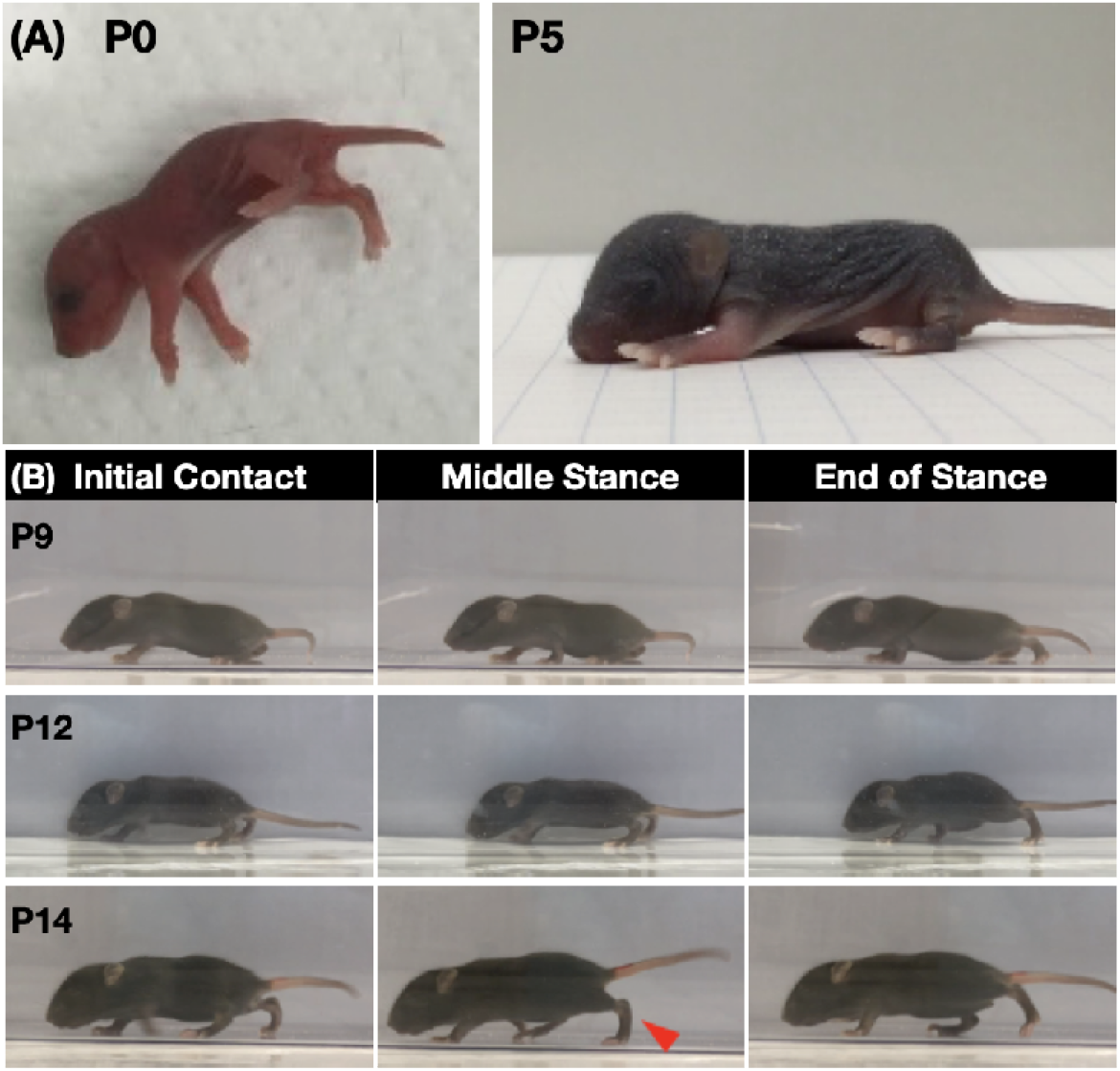
Summerly of limb movement and motor behavior in the postnatal phase. (A) P0 mouse limbs moved without weightbearing. Mice begin to bear weight at P5. (B) Progression of gait pattern with age. The red arrow shows the hock raise.

These are a few studies focusing on the mechanobiology of postnatal tendon formation compared with embryonic tendon development. It is difficult to define and quantify the applied mechanical force on the developmental tendons in the postnatal phase because their biomechanical situation changes dynamically. Two reports referred to the relationship between mechanical force and tendon development. Theodossiou et al. analyzed that mechanical development may proceed differently in weight-bearing Achilles tendons and non-weight-bearing tail tendons (S. K. Theodossiou et al., 2019). The same group then reported that the elastic modulus and cross-sectional area of the Achilles tendon were altered with the spinal cord transection (S. K.Theodossiou et al., 2021). They used a spinal cord transection rat model to prevent weight bearing but didn’t mention which affects the decreased muscle contraction or weight bearing. Finally, they suggested that the weight-bearing movement in the postnatal may provide the mechanical loading needed to direct postnatal tendon mechanical properties. We agree with their perspective on the importance of weight-bearing; moreover, we subdivided just weight-bearing and weight-bearing with muscle contraction from gait patterns. The results of the histological analysis in this study showed that the shape of the tendon cells nuclear was gradually elongated to the longitudinal direction of the tendon in the early postnatal phase (P0 to P7), and the number of tendon cells was reduced in the mid to late postnatal phase (P9 to P14). These results suggested that increasing the mechanical force may tune tendon cell turnover and extra-cellular matrix synthesis in this phase. Ansorge et al. focused on natural development and reported linear region stiffness and elastic modulus of Achilles tendons increased with age from P4 to P28. The distribution of fibril diameter sizes increased with age (Ansorge et al., 2011). Significant proliferation exists even as the tendon cells are reduced in density from P0 to P28 Achilles tendon (Grinstein et al., 2019). Summarizing these studies and our results, the postnatal tendon cell has a high sensitivity to the applied mechanical force, and cellular activities were induced by mechanobiological mechanisms. To understand how much mechanical force by the postnatal activity affects tendon cells, it is important to consider not only tendon growth but also tendon homeostasis and repair from injury. Our work is the first report to indicate the relationship between internal/external mechanical force and tendon development-specific gene expression in normal mice.

### 4.4. Limitation and Summary

This study is not without limitations. We used behavior tests to evaluate mechanical force, but we could not directly determine how much-increased muscle tension and changing muscle-tendon stretching occurred in this mouse model. This question should be addressed in future studies by measuring the actual muscle tensile strength in the triceps surae. Our results provide the baseline data and enable us to understand the part of the mechanotransduction mechanism more clearly through future studies using some transgenic mice. Finally, we detected the changing timeline in mechanical force for limbs using their physical behavior ability and patterns but could not test their frequency or cage activity. We have not yet answered the mechanism by which mouse embryos need muscle contraction. In the future, evaluating the extent to which their abilities are reflected in actual activities is necessary.

In summary, our findings underscore that tendon development needs mechanical force from changing limb movement depending on muscle contraction and external mechanical force (Fig. 8). External mechanical force, especially in the tendon maturation process, is a novel point of view for mechanotransduction in tendon development. Understanding the true mechanotransduction mechanism in tendon maturation provides insights into tendon degeneration and the treatment after tendon injury. Future studies need to focus on tendon development, not only the role of muscle contraction as an internal factor but also an external factor, such as the environment and gravity, between the embryo to the postnatal phase.

**Fig.8.**
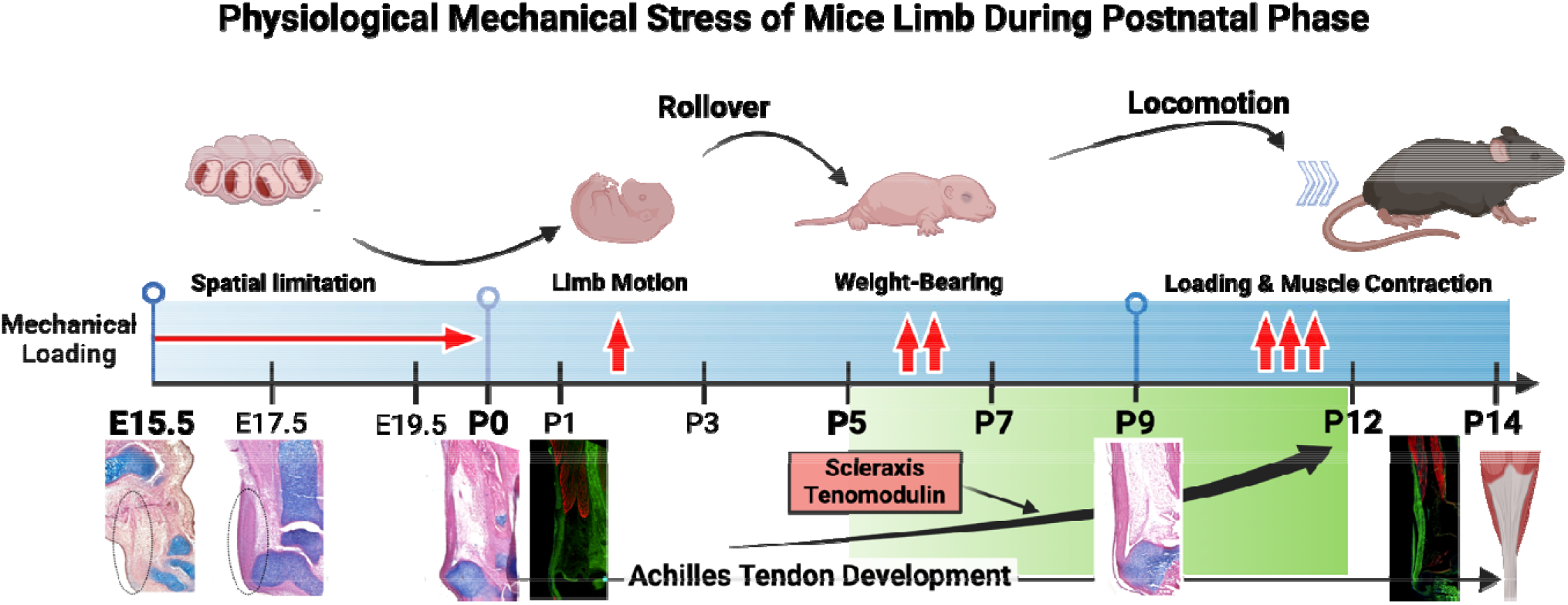
Summary of key findings during murine tendon development between embryo to early postnatal.

## CRediT authorship contribution statement

**Yuna Usami:** Writing – original draft, Visualization, Validation, Methodology, Investigation, Formal analysis, Data curation, Funding acquisition. **Xi Jiang:** Investigation, Methodology. **Nathaniel Dyment:** Writing – review & editing, Investigation, Methodology, Resources. **Takanori Kokubun:** Writing – review & editing, Supervision, Resources, Visualization, Project administration, Conceptualization.

## Data availability

The Data that supports the findings of this study are available in the supplementary material of this article.

## Acknowledgments

Graphical abstracts and Figures were created with BioRender.com.

